# Disconnects in global discourses—the unintended consequences of marine mammal protection on small-scale fishers

**DOI:** 10.1101/2020.01.01.892422

**Authors:** Katrina J. Davis, Joanna Alfaro-Shigueto, William N.S. Arlidge, Michael Burton, Jeffrey C. Mangel, Morena Mills, E.J. Milner-Gulland, José Palma Duque, Cristina Romero-de-Diego, Stefan Gelcich

**Affiliations:** Department of Zoology, University of Oxford, Zoology Research and Administration Building, 11a Mansfield Road, Oxford, OX1 3SZ, United Kingdom; UWA School of Agriculture & Environment, University of Western Australia, Stirling Highway, Crawley, Western Australia, Australia; Centre for Biodiversity and Conservation Science, The University of Queensland, St. Lucia, Queensland, Australia 4072; Pro Delphinus / Calle José Galvez 780E, Lima 15074, Perú; School of Biosciences, University of Exeter, Cornwall Campus, Penryn, Cornwall, TR10 9FE, United Kingdom; Facultad de Biología Marina, Universidad Científica del Sur, Ant Panamericana Sur km19, Lima 42, VES, Perú; Pembroke College, University of Oxford, St. Aldates, Oxford, OX1 1DW, United Kingdom; Faculty of Natural Sciences, Centre for Environmental Policy, Imperial College London, South Kensington Campus, London, SW7 2AZ, United Kingdom; Center of Applied Ecology and Sustainability, Pontificia Universidad Católica de Chile, Alameda 340, Santiago, Chile; School of Earth and Environmental Science, University of Queensland, St Lucia, Queensland, 4072, Australia

**Keywords:** Chile, human-wildlife conflict, marine mammals, Peru, pinnipeds, small-scale fisheries

## Abstract

Globally, the populations of many marine mammals remain of critical concern after centuries of exploitation and hunting. However, some marine mammal populations (e.g. pinnipeds) have largely recovered from exploitation, and interactions between these species and fisheries—particularly small-scale fisheries—is once again of concern globally. The large scope and widespread scale of interactions highlights the local disconnect between two global policies: marine mammal conservation and small-scale fisheries protection. In this research, we explore these conflicting global policies by assessing the perceptions of coastal small-scale fishers in Peru and Chile regarding their interactions with pinnipeds, including the South American sea lion (*Otaria flavescens*) and South American fur seal (*Arctocephalus australis*). We surveyed 301 gill net fishers and assess perceptions using a best-worst scaling methodology. We find that fishers are chiefly concerned with the increase in pinniped populations, perceive that their interactions with pinnipeds have significantly increased over the past 80 years, and report pinniped-driven catch and income losses ≥ 26 per cent. Surprisingly, fishers do not believe that compensation schemes will resolve this issue—instead they overwhelmingly call for pinniped population culls. The reported number of pinnipeds illegally killed by fishers suggests the potential for large negative impacts on these protected species, and a loss of legitimacy in marine regulation. Collectively, our results portray a sense of marginalisation from fishers’—that global policy treats them as less “important” than marine mammals. Our results highlight the increasing disconnect in global policy, which on one hand seeks to protect threatened marine mammal populations, and on the other seeks to promote the welfare of small-scale fishers.

## Introduction

Marine mammals perform fundamental roles in marine systems such as trophic regulation, nutrient cycling, and tourism opportunities (1–6). In the 19^th^ century, extreme collapses in global marine mammal populations occurred (7)—support for population control (through hunting or commercial extraction) was often driven by interactions with fisheries (8). These collapses eventually led to widespread protective legislation (9), including the International Whaling Convention (ca. 1940), and Marine Mammal Protection Acts (e.g. USA (1972), New Zealand (1978), UK (1981)). Following strict protection, many marine species (52%) are now recovering (7, 9), including most pinnipeds (e.g. seals, sea lions), though others (10%) are still declining (9). Parallel with the global focus on protecting marine mammal species, there has been an international push to recognise and protect the livelihoods of small-scale fishers. An estimated 22 ± 0.45 million people in the world are employed in the small-scale fishing sector, and they generally have very little income security (10). For these and other reasons, the Food and Agricultural Organization (FAO) has highlighted the importance of the small-scale fisheries sector as a target for improving income and food security in developing countries (11). Worryingly, increases in the abundance of some marine mammals are generating tension with fisheries, particularly small-scale coastal fisheries (12, 13). For example, in South America, pinniped depredation of catch is estimated to occur in ~56% of catches (14, 15), and lead to average economic losses of ~30 per cent (16). In extreme cases, fishers can react to conflict with mammals through illegal behaviour, which can result in shooting or poisoning them (17, 18). These actions ultimately exacerbate the human-wildlife conflict and threaten marine-mammal population recovery (19). The scale of interactions between small-scale fisheries and marine mammal populations, especially pinnipeds, is increasing. However, the local disconnect between these two global policies: marine mammal conservation and small-scale fisheries protection, has yet to be quantified and its nuances recognised in global fora. Furthermore, effective solutions remain unidentified.

Despite the potential for marine mammal–small-scale fisher interactions to lead to negative outcomes for both marine mammal populations and fisher welfare, this conflict remains poorly understood. This could be due to taboos regarding marine mammal killings (20), the potential for public outcry if these conflicts are publicised (16), and fisher’s fear of reprisals in the form of fines or sanctions from government agents. The wicked nature of the problem has meant that there are no straightforward solutions. This is partly due to concerns over affected social groups, e.g. small-scale fisher welfare. Nevertheless, homogenous application of marine mammal protection: which, in many locations does not discriminate between marine mammals that remain critically endangered (e.g. the vaquita, *Phocoena sinus*), and marine mammals whose populations are recovering (9) (e.g. pinnipeds such as sea lions and seals), is challenging the continued viability of some small-scale fisheries. The disconnect in these global discourses needs to be addressed and a coherent narrative that supports both conservation and small-scale fisheries developed.

Our aim in this research was to investigate the local impact of the increase in sea lion populations, which are a consequence of global conservation policies to protect marine mammals, on small-scale fishers. We base this research in the Eastern Pacific Rim, one of the largest upwelling systems in the world (21). Specifically, we focus on coastal areas spanning Peru and Chile (see Figure 1). Peru and Chile are the 5^th^ and 12^th^ biggest countries worldwide for fisheries landings (22), and their small-scale fisheries are highly integrated with world fisheries markets (23). These fisheries also overlap with pinniped foraging areas, increasing the potential for interactions between this sector and marine mammals (8, 12, 13). Peru’s small-scale fishing fleet contains an estimated 18,000 vessels and 67,000 fishers, distributed over 106 ports and landing sites (Castillo et al. 2018). The fleet has tripled in size since 1995. Landings in 2012 were in excess of 1 million tons, the majority of which was for human consumption in the Peruvian domestic market (24). In Chile, fisheries represent an important component of local economies and landings worldwide (25). During 2016, landings from Chilean small-scale fisheries were 1.7 million tons, over half of total fisheries landings (26). In our study area, there are two pinniped species: the South American sea lion (*O. flavescens*) and South American fur seal (*Arctocephalus australis*). Since the 1970’s, populations of South American sea lions (*O. flavescens*) in Peru and Chile have recovered strongly from commercial exploitation, which was banned fully in the 1990s. Conversely, since the 1997– 1998 ENSO event (19) fur seals are considered in danger of extinction along the Peruvian coast. Note, that in the material to follow we focus our discussion on the South American sea lion, as interactions are correspondingly more common in the study area (27). The case study area provides a unique opportunity to investigate the unintended consequences of marine mammal protection on small-scale fisheries. In this research, we quantify key motivations behind fishers’ struggles with sea lions, fishers’ perceptions regarding the impact of sea lions on catch and income, how interactions with sea lions have changed over time, and estimates of the number of animals illegally killed by fishers.

**Figure 1.**
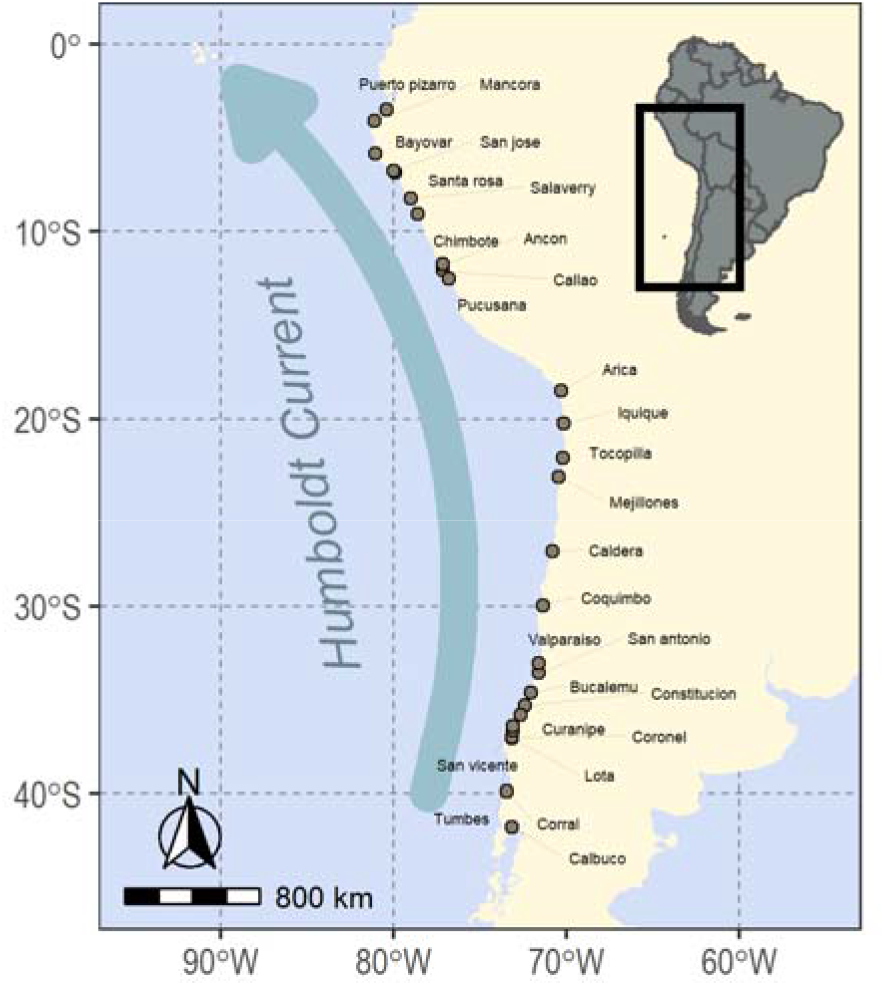
Survey locations along the coasts of Peru and Chile.

We surveyed 301 gillnet fishers in Chile (n = 201) and Peru (n = 100) using a best-worst scaling methodology to assess what aspect of their interactions with sea lions concerns them most. Best-worst scaling (BWS) is a form of discrete choice experiment, which asks respondents to choose the most important (best) or least important (worst) item from a list (28). The task is repeated a number of times, systematically varying the subset of items shown in each question. BWS is increasingly used in natural resource management applications as it facilitates evaluation of competing management alternatives (29). We developed a list of 12 reasons why sea lions might concern fishers based on the peer-reviewed literature and key informant interviews (Table 1). Note that we selected options to cover a range of possible economic, social, and ecological concerns. Because BWS asks respondents to rank options, options are not required to be mutually exclusive. The final survey was tested in 5 focus groups in Chile and Peru. We surveyed fishers in 17 locations in Chile, and 10 in Peru (Figure 1). We analysed responses using conditional logit (CL) (30) and scale-adjusted latent class (SALC) models (31), and calculated the marginal effects of preference class membership using a multinomial logit (MNL) model (32). We collected additional data on the impact of interactions on fishers’ catch and income, estimates of the number of sea lions killed by fishers, and how fishers perceive that interactions with sea lions have changed over time. We analysed changes in interactions over time using a double bounded tobit model (32).

**Table 1.** Reasons why sea lions could concern fishers.

| Coding | Full description |
| --- | --- |
| Strategy | Having to change my fishing strategy (e.g. location, gear, net placement) |
| Fish | Sea lions eat and scare fish from my nets |
| Inputs | Spending more money to repair damaged gear or travel further |
| Population | There are too many sea lions |
| Profits | Getting less money for damaged catch |
| Safety | Travelling further offshore to avoid sea lions endangers myself and my crew |
| Time | Working longer hours/Spending more time away from my family (e.g. repairing gear or longer fishing trips) |
| Behaviour | Sea lion behaviour is changing; they are no longer afraid of approaching fishing boats |
| Employment | Being forced to seek alternative employment |
| Reputation | Conflict with sea lions is giving fishing a bad reputation |
| Harm | Hurting sea lions while I am fishing |
| Risks | Sea lions may present unknown risks (e.g. disease) |

\*Note: English translation of survey, which was administered in Spanish.

## Results & Discussion

Approximately 30 per cent of the 301 gill net fishers that we surveyed in Peru and Chile were between 45 and 54 years of age, and on average had been fishing for 33 ± 14 years (mean ± SD). The first question regarding sea lions that fishers were asked was open-ended and asked respondents to describe the first word that came to their mind when they heard the term “sea lion”. A third of respondents responded “damage” or “losses” ^1^. Other responses included “bad”, “destruction”, and “rage.” A second open-ended question asked respondents to describe their interactions with sea lions. Nearly all (87 per cent) of the sample responded that their interactions are negative. These open-ended questions provide insight into how fishers frame their associations and concerns regarding sea lions (33).

We analysed what fishers thought was most concerning about their interactions with sea lions using conditional logit (CL) and scale adjusted latent class (SALC) models of best and worst responses. Results for both models are presented as importance scores, which describe the probability that a respondent will pick a given item as “best” from a set, assuming that all other items are of average importance. Results from the CL model indicated that fishers’ main concern is that sea lion populations are too large, thus increasing the probability for negative interactions (Figure 2.A). Using the SALC model we can investigate heterogeneity within the fisher population—whether there were groups of respondents with similar views, who might share similar characteristics (e.g. age, educational background) (Figure 2.B-E). The SALC analysis identified five groups of respondents who shared similar preferences, and two scale classes—groups who responded to questions with similar consistency (vs. inconsistency)—using effects coding (see Supplementary Material for model selection statistics).

**Figure 2.**
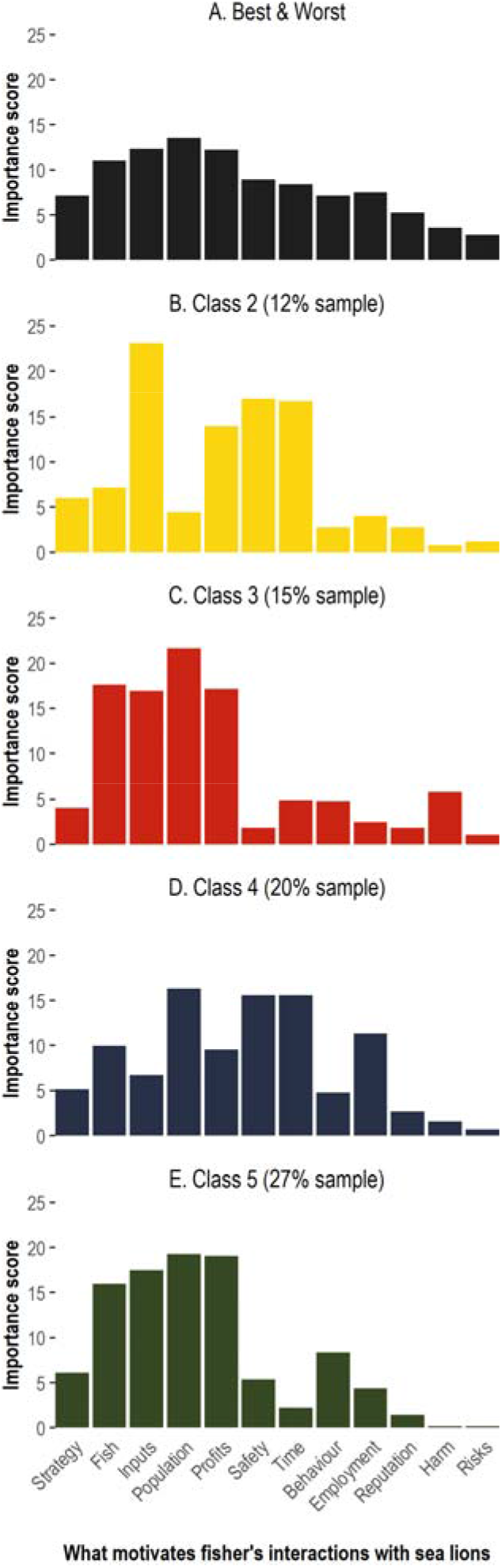
Importance scores for 12 reasons that sea lions may concern fishers in Peru and Chile. (A) Results from conditional logit model of best and worst responses, (B-E) scale adjusted latent class model of best and worst responses for preference class 2 to class 5 respectively. Class sizes are indicated in brackets. Note: Preference class 1 (~26 per cent of the sample) is not shown as this class has equal weights for all BWS options.

Class 5 (27% of sample) and Class 3 (15%) have a similar pattern of preferences for the most important concerns: sea lion population size, fish being eaten, increased input costs and reduced profits from damaged catch. Although sea lion population size is likely viewed as the underlying cause of the other impacts, this is not true for all classes: e.g. Class 2 (25%) have as their greatest concerns input costs, damage to catch, safety issues from increased travel and working longer hours. The sea lion population per se is not raised, but its consequences, and in particular a focus on changes in the working environment that are needed because of sea lions, are identified as chief concerns. Class 4 (20%) has a slightly more even allocation of concern, with again a concern about the time and safety issues. Of these four classes it is notable that there is little weight given to the impact on sea lions, or the implications of conflict for the reputation of the fishery. Class 1 (26%) displayed no differentiation across concerns. This may reflect that all 12 items are of equal importance, or (and we would hypothesis more likely) that these respondents were not engaging in the choice task but instead selecting best and worst at random. Note that the BWS approach identifies the relative concern surrounding the items within a class, but does not give any indication of absolute concern within or across classes.

To evaluate whether fishers’ characteristics could explain membership of preference classes we used a multinomial logit model. We found that involvement in sea lion tourism, perceived impact of sea lions on earnings, and nationality, significantly predicted preference class membership (see Supplementary Material for model results). This suggests that fishers’ perceptions of sea lion interactions are strongly linked with the impacts of sea lion on earnings, both negatively through depredation and positively through the provision of tourism opportunities. We analysed the marginal effects of these characteristics, which show how the probability of being in a given preference class changes when e.g. a respondent is involved in sea lion tourism. The probability of being in preference class 5—the largest class who identified increasing sea lion populations as their greatest concern—is 65 percentage points higher if the respondent is from Chile. Conversely, the probability of being in class 1—the class whose preferences are indistinguishable from random—is 57 percentage points higher if the respondent is from Peru. Fishers who think sea lions have a smaller negative impact on income (or greater positive impact) have a higher probability of being in class 1, and less probability of being in class 5. Put another way, every 10 percentage point decrease in the negative impact of sea lions on income relates to a 3.7 percentage point increase in the chance of being in class 1 and a 5.9 percentage point decrease in the chance of being in class 5. Respondents involved in sea lion tourism are between 20 and 16 percentage points more likely to be in preference class 3 and 4 respectively, and 26 percentage points less likely to be in class 1. Class 3 and 4 respondents were most concerned about increasing sea lion populations. Class 3 was also concerned about economic impacts, and Class 4 respondents were also concerned about safety and time spent away from family.

From our results, we identify that fishers’ chief concern is the need for control of sea lion populations—although this may underpin concerns about the economic impact that sea lions are having on their welfare. This result points to a disconnect between global marine mammal protection efforts, and international concerns over the welfare of small-scale fisheries. This suggests that unless fishers’ anxieties regarding the increase in marine mammal populations is addressed—global policies and regulations regarding marine mammal protection could lose legitimacy. We investigated whether this was already occurring in Peru and Chile by asking fishers to estimate the number of sea lions harmed by crew on an average vessel in their area. All questions were framed as impersonal, asking whether respondents had heard of other fishers engaging in the relevant activities, as harming sea lions is illegal in both Peru and Chile (34, 35). Note that not all fishers responded to these questions, hence the sample size is lower than the total number of survey respondents (n = 301). We first asked fishers if they had heard of other crew defending their catch from sea lions. Approximately 62 per cent of fishers (n = 298) said yes. We then asked fishers if they had heard of other crew killing sea lions to defend their catch. Approximately 69 per cent (n = 208) said yes. Finally, fishers (n = 126) estimated that crew on vessels in their area would kill on average 3 ± 9 (median ± SD) sea lions per month. Apart from the potential for this mortality to affect pinniped population viability, it is concerning because it implies a loss of legitimacy for marine regulations. This loss of legitimacy could spill over to other regulatory areas, and as a result, compliance rates with e.g., catch limits, could decrease. To add to perception of the illegitimacy of marine regulations, fishers also expressed a sense of marginalisation, of not being viewed as important as sea lions in marine policy. One fisher commented “I hope they do something, they protect the sea lions a lot.”^2^

### Perceptions of sea lion depredation and interactions over time

Fishers reported a general decrease in catch due to their most recent interactions with sea lions. Across the sample, 81 per cent of fishers reported catch losses of 26 per cent or higher. This equated to approximately a third of fishers reporting catch losses of either 26-50 per cent or 51-75 per cent and 24 per cent of fishers reporting catch losses between 76-100 per cent. The majority of respondents (71 per cent) identified that, on average, the impact of sea lion interactions on their take-home income is a decrease of 41 per cent or greater. No significant difference was identified between fishers from Peru and Chile. These results have significant implications for the welfare of small-scale fishers in Peru and Chile—the majority of which are entry-level fishers with very small profit margins (36). Fishers were asked to assess the level of interaction with sea lions when they first started fishing (number of trips with interactions per 10 trips). Given the time period for when they first started fishing was from 1944 to 2018, this gives us an opportunity to investigate the (subjective) trend in interactions over time. To evaluate how fishers perceive their interactions with sea lions have changed over time, we used a double bounded tobit model. Results show that fisher’s interactions with sea lions have increased by 10 per cent every decade (p-value < 0.001). These results can help explain fishers’ preoccupation with sea lion population numbers, which have led to increasing numbers of interactions throughout the period a fisher has been fishing. Currently, fishers are estimated to have interactions with sea lions in 9 fishing trips out of 10 (intercept = 9.3, p-value <0.001).

We asked fishers a number of additional questions to understand the role that fishing plays in their lives, along both economic and social dimensions. Over 77 per cent of fishers obtain between 76 and 100 per cent of their income from fishing, and 85 per cent are the chief income earners in their household. The overwhelming majority of fishers (74 per cent) strongly agreed that being a fisher is very important to them. Slightly less than half of the sample responded that they would be very uncomfortable working in another industry. These responses suggest that fishers depend on fishing, not just as their main form of income, but also for their identity. The dependence of respondents on fishing highlights how vulnerable fishers and their households are to the large negative impacts that sea lion interactions have on catch and incomes.

We compare responses to a further attitudinal question: “I consider myself environmentally friendly” with fisher’s agreement with the following statement, “sea lions are a pest.” This last question was coded on a Likert scale, with ‘1’ equal to ‘strongly disagree’ and 10 equal to ‘strongly agree’. Over 65 per cent of the sample described themselves as environmentally friendly *and* agreed that sea lions are a pest (scores greater than 8 on both questions). Next, we assessed whether this “pest score” could be explained by any other variables using a backward stepwise algorithm in R (MASS package (37)), using AIC model selection criteria. We found that fishers’ who would be more comfortable working outside of fishing or who were from Peru were significantly less likely to view sea lions as a pest (p < 0.05). This is logically consistent, as sea lions pose a greater risk to fishers who are more dependent on fishing. Respondents who listed a greater number of recent interactions with sea lions were significantly more likely to think sea lions were a pest (p < 0.01).

### How fishers frame solutions

Popular management actions to manage human–wildlife conflict include attempts to separate problem animals from affected human populations, for example, fencing to keep elephants from crops (38); compensation (39); use of equipment/infrastructure to control interactions e.g. devices to deter sea lions (15); and lethal control—either of problem animals (40) or of the larger population (41). We asked fishers what they thought would be the best solution to manage their interactions with sea lions. Fishers in our study area (~72 per cent of respondents) overwhelming support sea lion population control, through culls or regulated harvesting, as the best way to manage their interactions with sea lions. Compensation and devices to deter sea lions are suggested by only 5 per cent of fishers. This result mirrors findings from the BWS questions (Figure 2)—that the sheer number of sea lions is fishers’ greatest concern. Approximately 11 per cent of the sample report that there is no solution. This could be interpreted as fishers expressing their feelings of disempowerment, furthermore, that conflict with sea lions is not their responsibility to fix (e.g. it’s the government’s responsibility). One fisher suggested that the answer lies in reducing pressure on fish stocks (particularly Peruvian anchoveta) so that sea lions would have more food. Surprisingly, only a few fishers identified compensation or other forms of help as the best way of resolving conflict with sea lions. This may be because fishers have low familiarity with compensation schemes, suggesting that existing solutions to marine mammal conflict with fisheries is path dependent—we continue to manage conflict as it has been managed for ~200 years (8, 34), either through culling marine mammal populations (e.g., fishers win), or total protection of these species (e.g., fishers loose). Responses can be grouped into three broad categories: *separating* the problem, for example, reserve creation; *removing* the problem, through population culls; and *living* with the problem, for example through providing compensation for damaged catch or changing fishing practices.

Finally, we asked fishers who they thought was responsible for managing interactions with sea lions (Table 4). The majority of fishers felt the government was responsible for managing conflict with sea lions.

**Table 2.** Marginal effects describing fisher’s probability of being in each of the five preference classes based on their characteristics.

| Variable | Class 1 | Class 2 | Class 3 | Class 4 | Class 5 |
| --- | --- | --- | --- | --- | --- |
| Involved in sea lion tourism | -0.257 | -0.023 | 0.192 | 0.163 | -0.074 |
| Impact of sea lions on earnings | 0.037 | 0.010 | 0.002 | 0.009 | -0.059 |
| Respondents from Peru | 0.566 | 0.149 | -0.017 | -0.053 | -0.644 |

Notes: Membership of preference classes was modelled probabilistically as a function of individual specific characteristics using a multinomial logit functional form.

**Table 3.** Variables predicting response to agreement with “Sea lions are a pest.”

| Variable | Coefficient |  |
| --- | --- | --- |
| (Intercept) | 6.52 (0.86) | *** |
| Proportion of income from fishing | 0.27 (0.17) |  |
| Involved in sea lion tourism | 0.37 (0.28) |  |
| Comfortable working outside of fishing | -0.08 (0.04) | ** |
| Recent number of sea lion interactions | 0.25 (0.06) | *** |
| Respondents from Peru | -1.75 (0.25) | *** |

Note: Standard errors are shown in italics in brackets. \*\*,\*\*\* indicate p-values < 0.05, 0.001.

**Table 4.** Solutions and the actors that fishers perceive are responsible for managing interactions with sea lions.

| Solution:<br>sub-category | % | Solution:<br>broad-category | % | Responsible for<br>management | No. of<br>fishers |
| --- | --- | --- | --- | --- | --- |
| Population control | 60.1 | Remove | 71.8 | Government | 240 |
| Exploit | 11.6 |  |  | Military | 34 |
| Reserve | 3.3 | Separate | 3.3 | NGOs | 15 |
| No solution | 10.6 | Live with | 23.3 | Industry/fishers | 11 |
| Change fishing practice | 4.7 |  |  | Don’t know | 11 |
| Technology | 4.0 |  |  | Scientific bodies | 4 |
| Research | 2.7 |  |  | Other | 6 |
| Compensation/help | 1.3 |  |  |  |  |
| Change the law | 0.7 | NA |  |  |  |

Note: In the final column (“No. of fishers”), fishers could suggest more than one agency is responsible for management, hence responses do not sum to 301 (the number of respondents).

Global policy is attempting to support small-scale fisheries. This includes through directives from the Food and Agricultural Organisation (11), shifts to co-management systems (42), and a focus on incorporating local knowledge into management decisions (43). Here, we demonstrate that small-scale fishers’ perceptions and experiences regarding the impacts of marine mammals are *not* reflected in global policies prioritising marine mammal conservation. Among a range of problems they could have with pinnipeds, fishers’ chief concern regards the size of pinniped populations; they also perceive that their interactions with these animals have been significantly increasing over time. Fishers further report large negative impacts on catch and income as a result of interactions with pinnipeds, and large numbers of pinnipeds illegally killed in retaliation. The key message from these results is that it is often local communities who are bear the costs of marine conservation success (40, 44).

In the marine realm, conflict between marine mammals and fisheries continues to be managed as it has been for the past ~200 yrs (8, 34)—either by culling marine mammal populations to prevent negative impacts on fisheries, or by enacting broad scale protection for mammals—with little concession made for species’ current threat status. Compared to experiences in the marine realm, terrestrial human–wildlife conflict management has taken a more nuanced approach, with some lessons applicable to marine settings—particularly strategies focusing on stakeholder engagement. These include insurance mechanisms, incentives, or compensation (40). Notwithstanding, there are important challenges that managing conflict in marine settings imposes on managers that are not experienced in terrestrial settings, such as those inherent to physically operating in the ocean, and the greater spatial distribution and mobility of marine species (40).

To avoid alienating fishers, and hence loss of legitimacy in marine policy, the nuances associated with interactions between marine mammals and small-scale fishers need to be addressed in global fora. This implies the need for open dialogue with fishers, and avoiding broadly enacted conservation policy that treats all marine mammals equivalently. By incorporating the needs and opinions of fishers in global dialogue, marine mammal policies have more chance of finding solutions that allow small-scale fisheries to continue operations while maintaining viable marine mammal populations. Furthermore, if fishers remain engaged with policy development, so that they feel protected and heard, this should help maintain and improve the legitimacy of broader marine regulation.

## Methods

### Survey design

Our survey featured four main sections focusing on: (1) fishers’ fishing activities; (2) fisher’s motivations for concern regarding their interactions with pinnipeds (best-worst scaling questions); (3) fishers’ interactions with pinnipeds; and (4) fishers’ socio-demographic information. The survey was administered in Spanish using face-to-face questionnaire-based interviews by a research team in coastal areas in Peru and Chile. The survey was administered in 10 locations in Peru, and 17 locations in Chile (see Figure 1). We selected sampling locations in Peru to capture major gillnet fishery sites with representation in each geopolitical region to ensure geographical coverage. Sampling locations in Chile were selected to include *caletas* (fishing coves) with the highest landings of pelagic and demersal fish species, and to ensure equal coverage across the south, central and northern regions of Chile.

To select attributes for the best-worst scaling survey, we began with a brainstorm of potential concerns regarding interactions between fishers and pinnipeds. These were grouped into six themes: ecological, economic, social, fishing-approach/method/flexibility, and capacity-related. We began with a list of 20 options, which were narrowed down to 12 (see Table 1) through key informant interviews and following focus groups in Peru and Chile. Our best-worst scaling design had 12 choice sets with 5 options seen per choice set, and 8 survey versions. We assumed sequential best–worst ranking. Choice sets were designed using Sawtooth software (45). The order in which survey versions were administered was randomised to ensure equal version coverage.

### Modelling approach

We assessed fishers chief concern regarding their interactions with pinnipeds using best-worst scaling (BWS), a form of discrete choice experiment. Discrete choice experiments (46) contain a number of choice sets which require the respondent to choose their “best” option from varying sets of three or more options (47). In the case of BWS, a “worst” option is also chosen; the difference between best and worst choices is assumed to encompass the largest perceptual difference on an underlying continuum of interest for the respondent (47). Analysis of choices allows each option to be rank-ordered on a common scale and assessed on the basis of its relative importance (48). We used a ‘case 1’ BWS—this case does not differentiate options according to attributes and is used to assess simple concepts (49). Following convention, our analytical approach is based on Random Utility Theory (50).

We employed a conditional logit model to evaluate the sample average, this model pools best and worst responses and the interested reader is directed to Louviere, Flynn and Marley (28) for a description of the model employed. To assess heterogeneity in perceptions across the sample, we drew on the scale adjust latent class model, following Rigby, Burton and Lusk (31). Membership of scale and preference classes was assessed using a multinomial logit model (32). Finally, to assess how respondents viewed their interactions with pinnipeds had changed over time, we employed a double bounded tobit model (32). Data were cleaned and summary statistics run in R (51). All logit and tobit models were analysed in Stata 15 (52), and latent class models in LatentGold (53).

## Acknowledgements

The authors would like to thank C. Ortiz, E. Alfaro, A. Jimenez, S. Pingo, A. Pasara, and E. Campbell for their assistance with surveys. The authors also acknowledge the support of Reserva Nacional de Islas e Islotes and Puntas Guaneras of SERNANP. Ethics approval [eUEBS000824 v2.0] was obtained prior to interviews.

## Footnotes

1 In Spanish: “daño”, dañino”, “perdidas”, or “malogra/perjudica.”

2 In Spanish: “Ojalá que se haga algo se protégé mucho a los lobos”

