## Supplementary Material for "Disconnects in global discourses—the unintended consequences of marine mammal protection on small-scale fishers"

In this section, we present a range of supplementary material for the article “Disconnects in global discourses—the unintended consequences of marine mammal protection on small-scale fishers.” Below, we present the test statistics for model selection among the evaluated models.

Table 1. Model selection criteria for scale-adjusted latent class models describing fishers’ chief concerns regarding their interactions with pinnipeds in Peru and Chile .

|  |  | **LL** | **BIC(LL)** | **CAIC(LL)** | **Npar** | **L²** | **df** | **p-value** | **Class.Err.** | **R²(0)** | **R²** |
| --- | --- | --- | --- | --- | --- | --- | --- | --- | --- | --- | --- |
| Model1 | 1-Class Choice | -9798.4135 | 19659.5319 | 19670.5319 | 11 | 19596.827 | 288 | 5.7e-3933 | 0 | 0.0784 | 0.084 |
| Model2 | 2-Class Choice | -9450.3116 | 19037.4338 | 19061.4338 | 24 | 18900.6231 | 275 | 6.4e-3796 | 0.0493 | 0.1263 | 0.1317 |
| Model3 | 3-Class Choice | -9312.8807 | 18836.6779 | 18873.6779 | 37 | 18625.7615 | 262 | 4.4e-3749 | 0.071 | 0.1481 | 0.1533 |
| Model4 | 4-Class Choice | -9246.0217 | 18777.0656 | 18827.0656 | 50 | 18492.0434 | 249 | 1.5e-3732 | 0.0837 | 0.1613 | 0.1665 |
| Model5 | 5-Class Choice | -9187.4372 | 18734.0024 | 18797.0024 | 63 | 18374.8744 | 236 | 1.1e-3719 | 0.0797 | 0.1715 | 0.1767 |
| Model6 | 6-Class Choice | -9145.8805 | 18724.9947 | 18800.9947 | 76 | 18291.761 | 223 | 3.1e-3714 | 0.0739 | 0.1783 | 0.1835 |
| Model7 | 7-Class Choice | -9110.2788 | 18727.897 | 18816.897 | 89 | 18220.5575 | 210 | 1.7e-3711 | 0.071 | 0.1872 | 0.1923 |
| Model8 | 2-sClass 1-Class Choice | -9497.1257 | 19068.3572 | 19081.3572 | 13 | 18994.2514 | 286 | 7.0e-3806 | 0 | 0.1253 | 0.1307 |
| Model9 | 2-sClass 2-Class Choice | -9342.0417 | 18832.295 | 18858.295 | 26 | 18684.0835 | 273 | 2.0e-3751 | 0.1207 | 0.1492 | 0.1544 |
| Model10 | 2-sClass 3-Class Choice | -9271.484 | 18765.2854 | 18804.2854 | 39 | 18542.9681 | 260 | 3.3e-3733 | 0.0864 | 0.1593 | 0.1645 |
| Model11 | 2-sClass 4-Class Choice | -9216.12 | 18728.6631 | 18780.6631 | 52 | 18432.24 | 247 | 1.3e-3721 | 0.109 | 0.1707 | 0.1759 |
| Model12 | 2-sClass 5-Class Choice | -9160.5353 | 18691.5995 | 18756.5995 | 65 | 18321.0707 | 234 | 4.9e-3710 | 0.1152 | 0.1819 | 0.187 |
| Model13 | 2-sClass 6-Class Choice | -9120.3115 | 18685.2575 | 18763.2575 | 78 | 18240.6229 | 221 | 3.5e-3705 | 0.1047 | 0.1878 | 0.1929 |
| Model14 | 2-sClass 7-Class Choice | -9087.5038 | 18693.748 | 18784.748 | 91 | 18175.0076 | 208 | 1.2e-3703 | 0.1497 | 0.1956 | 0.2006 |
| Model15 | 2-sClass 2-Class Choice | -9439.99 | 18965.4867 | 18980.4867 | 15 | 18879.9801 | 284 | 2.9e-3783 | 0.0552 | 0.1341 | 0.1394 |
| Model16 | 2-sClass 3-Class Choice | -9289.2777 | 18738.1678 | 18766.1678 | 28 | 18578.5554 | 271 | 1.1e-3730 | 0.098 | 0.1575 | 0.1627 |
| Model17 | 2-sClass 4-Class Choice | -9225.9619 | 18685.642 | 18726.642 | 41 | 18451.9238 | 258 | 1.4e-3715 | 0.109 | 0.1697 | 0.1749 |
| Model18 | 2-sClass 5-Class Choice | -9171.0762 | 18649.9763 | 18703.9763 | 54 | 18342.1524 | 245 | 3.5e-3704 | 0.1199 | 0.1809 | 0.1861 |
| Model19 | 2-sClass 6-Class Choice | -9128.7737 | 18639.4771 | 18706.4771 | 67 | 18257.5474 | 232 | 2.6e-3698 | 0.1154 | 0.1876 | 0.1927 |
| Model20 | 2-sClass 7-Class Choice | -9094.3954 | 18644.8263 | 18724.8263 | 80 | 18188.7909 | 219 | 5.6e-3696 | 0.1538 | 0.1934 | 0.1985 |

We then present the full model results for this scale adjusted latent class model (Table 2), including coefficients for scale and preference classes. Finally, we present the results of a multinomial logit model explaining preference and scale class membership (Table 3).

Table 2. Scale-adjusted latent class model describing respondent’s choices.

| **term** |  |  | **coef** | **s.e.** | **z-value** | **p-value** |  |  |  |  |
| --- | --- | --- | --- | --- | --- | --- | --- | --- | --- | --- |
| sclass(1) | 1 |  | 0 | . | . | . |  |  |  |  |
| sclass(2) | 1 |  | 8.108 | 2.8266 | 2.8685 | 0.0041 |  |  |  |  |
| sclass(1) | seguro |  | 0 | . | . | . |  |  |  |  |
| sclass(2) | seguro |  | -1.0489 | 0.3803 | -2.7579 | 0.0058 |  |  |  |  |
| Attributes | Class 1 | z-value | Class 2 | z-value | Class 3 | z-value | Class 4 | z-value | Class 5 | z-value |
| Change fishing strategy | 0 | . | -0.032 | -0.140 | -0.512 | -2.542 | -0.299 | -1.775 | 0.217 | 1.323 |
| Sea lions eat & scare fish from nets | 0 | . | 0.182 | 0.757 | 1.658 | 5.942 | 0.558 | 2.997 | 1.868 | 11.193 |
| Spending on damaged nets/travelling further | 0 | . | 2.467 | 8.453 | 1.564 | 6.693 | 0.030 | 0.179 | 2.126 | 11.439 |
| Too many sea lions | 0 | . | -0.397 | -1.458 | 2.222 | 7.381 | 1.374 | 6.580 | 2.469 | 13.452 |
| Less money for damaged catch | 0 | . | 1.191 | 4.522 | 1.593 | 5.549 | 0.499 | 2.886 | 2.426 | 12.216 |
| Travelling further is dangerous | 0 | . | 1.590 | 6.905 | -1.393 | -6.021 | 1.288 | 5.868 | 0.050 | 0.261 |
| Longer hours/away from family | 0 | . | 1.552 | 6.073 | -0.293 | -1.118 | 1.287 | 5.931 | -0.974 | -5.616 |
| Sea lion behaviour is changing | 0 | . | -0.939 | -4.257 | -0.324 | -1.454 | -0.384 | -2.245 | 0.648 | 3.477 |
| Forced to seek alternative employment | 0 | . | -0.516 | -1.826 | -1.074 | -5.063 | 0.750 | 4.162 | -0.200 | -1.133 |
| Conflict gives fishers bad reputation | 0 | . | -0.938 | -3.988 | -1.397 | -5.374 | -1.026 | -4.494 | -1.445 | -8.195 |
| Hurting sea lions while fishing | 0 | . | -2.316 | -7.579 | -0.078 | -0.304 | -1.594 | -6.030 | -3.466 | -11.590 |
| Sea lions may present unknown risks |  |  |  |  |  |  |  |  |  |  |

Table 3. Multinomial logit model explaining preference and scale class membership, class sizes are also shown.

|  | Class 1 |  | Class 2 |  | Class 3 |  | Class 4 |  | Class 5 |  |
| --- | --- | --- | --- | --- | --- | --- | --- | --- | --- | --- |
| Socio-economic variable | coef | z-value | coef | z-value | coef | z-value | coef | z-value | coef | z-value |
| 1 | 0.433 | 1.129 | -0.121 | -0.271 | -0.237 | -0.582 | 0.450 | 1.220 | -0.524 | -1.282 |
| Peru | 2.060 | 5.879 | 1.094 | 2.735 | -0.230 | -0.500 | -0.376 | -0.925 | -2.548 | -3.545 |
| Involved in sea lion tourism | -1.116 | -2.105 | -0.318 | -0.560 | 1.175 | 3.203 | 0.665 | 1.750 | -0.407 | -0.923 |
| Impact of sea lions on earnings | 0.131 | 2.221 | 0.072 | 0.982 | 0.002 | 0.039 | 0.029 | 0.502 | -0.234 | -3.864 |
| Class size |  |  |  |  |  |  |  |  |  |  |
| Preference | 1 |  | 2 |  | 3 |  | 4 |  | 5 |  |
|  | 0.260 |  | 0.123 |  | 0.147 |  | 0.205 |  | 0.265 |  |
| Scale | 1 | 2 |  |  |  |  |  |  |  |  |
|  | 0.487 | 0.513 |  |  |  |  |  |  |  |  |

Below, we include a table with the marginal effects of changes in each of the four socio-demographic variables that explain preference class membership.

Table 4. Marginal effects of changes in socio-demographic variables explaining preference class membership.

| Variable | Class 1 | Class 2 | Class 3 | Class 4 | Class 5 |
| --- | --- | --- | --- | --- | --- |
| 1 | 0.108 | -0.017 | -0.037 | 0.089 | -0.143 |
| Peru | 0.566 | 0.149 | -0.017 | -0.053 | -0.644 |
| Involved in sea lion tourism | -0.257 | -0.023 | 0.192 | 0.163 | -0.074 |
| Impact of sea lions on earnings | 0.037 | 0.010 | 0.002 | 0.009 | -0.059 |
